# Diminished Cortical Thickness is Associated with Impulsive Choice in Adolescence

**DOI:** 10.1101/230888

**Authors:** Marieta Pehlivanova, Daniel H. Wolf, Aristeidis Sotiras, Antonia Kaczkurkin, Tyler M. Moore, Rastko Ciric, Philip A. Cook, Angel Garcia de La Garza, Adon Rosen, Kosha Ruparel, Anup Sharma, Russell T. Shinohara, David R. Roalf, Ruben C. Gur, Christos Davatzikos, Raquel E. Gur, Joseph W. Kable, Theodore D Satterthwaite

## Abstract

Adolescence is characterized by both maturation of brain structure and increased risk of negative outcomes from behaviors associated with impulsive decision-making. One important index of impulsive choice is delay discounting (DD), which measures the tendency to prefer smaller rewards available soon over larger rewards delivered after a delay. However, it remains largely unknown how individual differences in structural brain development may be associated with impulsive choice during adolescence. Leveraging a unique large sample of 427 human youths (208 males and 219 females) imaged as part of the Philadelphia Neurodevelopmental Cohort, we examined associations between delay discounting and cortical thickness within structural covariance networks. These structural networks were derived using non-negative matrix factorization, an advanced multivariate technique for dimensionality reduction, and analyzed using generalized additive models with penalized splines to capture both linear and nonlinear developmental effects. We found that impulsive choice, as measured by greater discounting, was most strongly associated with diminished cortical thickness in structural brain networks that encompassed the ventromedial prefrontal cortex, orbitofrontal cortex, temporal pole, and temporoparietal junction. Furthermore, structural brain networks predicted DD above and beyond cognitive performance. Taken together, these results suggest that reduced cortical thickness in regions known to be involved in value-based decision-making is a marker of impulsive choice during the critical period of adolescence.

**SIGNIFICANCE:** Risky behaviors during adolescence, such as initiation of substance use or reckless driving, are a major source of morbidity and mortality. In this study, we present evidence from a large sample of youths that diminished cortical thickness in specific structural brain networks is associated with impulsive choice. Notably, the strongest association between impulsive choice and brain structure was seen in regions implicated in value-based decision-making; namely, the ventromedial prefrontal and orbitofrontal cortices. Moving forward, such neuroanatomical markers of impulsivity may aid in the development of personalized interventions targeted to reduce risk of negative outcomes resulting from impulsivity during adolescence.

## INTRODUCTION

Adolescence is marked by an increased vulnerability to risky behaviors, such as tobacco, alcohol, and drug use, reckless driving, and unprotected sex (Eaton et al., 2011). During this vulnerable period, the brain undergoes dramatic structural changes (Giedd et al., 1999; Sowell et al., 2004). Some evidence suggests that risk during adolescence is associated with differential maturation of brain regions related to reward processing (such as the orbitofrontal cortex and ventral striatum) and those necessary for cognitive control (such as the dorsolateral prefrontal cortex, dlPFC; Casey et al., 2008; Van Leijenhorst et al., 2010). One of the most commonly used indices of impulsive choice is delay discounting (DD)— a behavioral measure of impulsivity where one chooses between a smaller reward delivered sooner, and a larger reward with a longer delay (Kirby and Maraković, 1995; Peters and Büchel, 2011; Kable, 2013). Delay discounting engages regions known to mature at different rates in adolescence, including dlPFC (Peters and Büchel, 2011), orbitofrontal cortex, and ventral striatum (Kable and Glimcher, 2007; Bartra et al., 2013). Increased DD has been proposed as a framework for understanding substance abuse and other risky decisions as reflecting impulsive choices of immediate rewards (Bickel et al., 2007). Indeed, studies of adolescents show that higher impulsivity, as indexed by higher discounting, is associated with increased smoking frequency (Reynolds, 2004), greater alcohol consumption (Field et al., 2007), and predicts longitudinal increase in both smoking (Audrain-McGovern et al., 2009) and alcohol use (Fernie et al., 2013).

At present, it remains relatively unknown how individual differences in structural brain development may relate to DD in adolescents. Neuroanatomical studies in adults are more numerous, but have yielded inconsistent results, perhaps due to small samples and focused region-of-interest analyses (for a review see Kable and Levy, 2015). For example, it has been reported that greater DD (more impulsive choice) is associated with reduced gray matter volume in lateral prefrontal cortex (Bjork et al., 2009), superior frontal gyrus (Schwartz et al., 2010), and putamen (Dombrovski et al., 2012; Cho et al., 2013). Furthermore, greater DD has been associated with larger volume of the ventral striatum and posterior cingulate cortex (PCC, Schwartz et al., 2010), medial prefrontal regions and anterior cingulate cortex (ACC, Cho et al., 2013), as well as prefrontal cortex (Wang et al., 2016). One study of cortical thickness (CT) in adults revealed an association between higher DD and decreased CT in both medial prefrontal cortex and the ACC (Bernhardt et al., 2014). To our knowledge, there have been no neuroanatomical studies in adolescents that specifically examine the relationship between DD and cortical thickness. Notably, findings from adults may not necessarily extend to adolescents, given the dynamic re-modeling of brain structure that occurs during this critical period (Sowell et al., 2004).

Accordingly, here we investigated how individual differences in DD may be associated with differences in brain structure during adolescence. To do this, we capitalized on a large sample of 427 youths imaged as part of the Philadelphia Neurodevelopmental Cohort (Satterthwaite et al., 2014a; 2016). We delineated covariance networks of cortical thickness using a recently-developed application of nonnegative matrix factorization for the multivariate analysis of high-dimensional neuroimaging data (Sotiras et al., 2015; 2017). We evaluated the association between DD and CT in each network, while specifically modeling both linear and nonlinear developmental effects using penalized splines. We hypothesized that we would find associations between DD and CT in brain regions associated with reward processing, such as the ventromedial prefrontal cortex (vmPFC; Kable and Glimcher, 2007; Bartra et al., 2013), as well as regions subserving cognitive control (e.g. dlPFC). As described below, diminished CT in these as well as other networks was associated with impulsive choice, and predicted individual variation in DD above and beyond that explained by cognitive performance.

## MATERIALS AND METHODS

### Participants and sample construction

Participants were a subsample of 1,601 youths recruited as part of the Philadelphia Neurodevelopmental Cohort (PNC) who underwent neurocognitive assessment (Gur et al., 2010; 2012), as well as neuroimaging (Satterthwaite et al., 2014a; 2016). A sub-sample of PNC participants (*n* = 453) completed the delay discounting (DD) task. Of those, *n* = 2 did not pass the quality control criteria for the task (described below). Additionally, *n* = 24 participants were excluded for the following reasons: health conditions that could impact brain structure (*n* = 19), scanning performed more than 12 months from DD testing (*n* = 1), inadequate structural image quality (*n* = 3) or missing imaging data (*n* = 1). The remaining *n* = 427 participants constituted our final sample for analysis (mean age at scanning: 17.0 ± 3.2 years, age range: 9.3–24.3 years; 48.7%, *n* = 208 males).

### Delay discounting task

The DD task consisted of 34 self-paced questions where the participant chose between a smaller amount of money available immediately or a larger amount available after a delay. This task was modeled after the work of Senecal et al. (2012). The smaller, immediate rewards ranged between $10 and $34 and were always displayed at the top of the computer screen. The larger, delayed rewards were fixed at $25, $30, or $35, with the delays ranging between 1 and 171 days. Larger, delayed rewards were always displayed on the bottom of the screen. All rewards were hypothetical but participants were instructed to make decisions as if the choices were real. Discount rates based on hypothetical choices have shown no systematic differences from discount rates based on real rewards, in the same participants (Johnson and Bickel, 2002). The set of choices was identical in content and order for all participants. The DD task was administered as part of an hour-long web-based battery of neurocognitive tests (Computerized Neurocognitive Battery, described below), on a separate day from the imaging session. The mean interval between the DD task and imaging was 0.44 months with a SD of 1 month (range 0–8 months).

Discount rates from the delay discounting task were calculated assuming a hyperbolic discounting model of the form: *SV* = *A*/(1 +*kD*), where *SV* is the subjective value of the delayed reward, *A* is amount of the delayed reward, *D* is the delay in days, and *k* is the subject-specific discount rate (Mazur, 1987). We used the fmincon optimization algorithm in MATLAB (Mathworks, Natick, MA; RRID:SCR_001622) to estimate the best-fitting *k* from each participant’s choice data. A higher *k* value indicates steeper discounting of delayed rewards and thus more impulsive choices. As the distribution of discount rates is highly right-skewed, we used log-transformed *k* (log *k*) in all analyses.

We performed quality control to ensure that participants were not responding randomly, and verified that their responses were a function of task variables which should be relevant to the choice. Although a hyperbolic discounting model has been shown to fit discounting data better than an exponential model (Kirby and Maraković, 1995), quality control was performed independently of assumptions about the shape of the discount function. Specifically, each participant’s responses were fit using a logistic regression model, with predictors including the immediate amount, delayed amount, delay, their respective squared terms, and two-way interaction terms. We assessed goodness of fit of this model using the coefficient of discrimination (Tjur, 2009), and discarded DD data from any participant who had a value of less than 0.20.

### Neurocognitive battery

Cognition was assessed using the University of Pennsylvania Computerized Neurocognitive Battery (Penn CNB, Gur et al., 2010; 2012) during the same session that delay discounting was evaluated. Briefly, this hour-long battery consisted of 14 tests administered in a fixed order, evaluating aspects of cognition, including executive control, episodic memory, complex reasoning, social cognition, and sensorimotor and motor speed. Except for the motor tests that only measure speed, each test provides measures of both accuracy and speed. Performance on the tests for each domain is summarized as cognitive factors obtained with exploratory factor analysis with an oblique rotation (Moore et al., 2015). Prior work has demonstrated that accuracy on this battery can be parsimoniously summarized as either one overall cognitive performance factor or three domain-specific factors, including executive function and complex reasoning combined, social cognition, and episodic memory (Moore et al., 2015). Associations between DD and factor scores for each of these dimensions were analyzed, as described below.

### Image acquisition and quality assurance

Image acquisition and processing are reported in detail elsewhere (Satterthwaite et al., 2014a; 2016). Briefly, all data were acquired on a single scanner (Siemens TIM Trio 3 Tesla, Erlangen, Germany; 32-channel head coil) using the same imaging sequences for all participants. Structural brain scanning was completed using a magnetization-prepared, rapid acquisition gradient-echo (MPRAGE) T1-weighted image with the following parameters: TR 1810 ms; TE 3.51 ms; FOV 180×240 mm; matrix 192×256; 160 slices; slice thickness/gap 1/0 mm; TI 1100 ms; flip angle 9 degrees; effective voxel resolution of 0.93 × 0.93 × 1.00 mm; total acquisition time 3:28 min.

### Image quality assurance

T1 image quality was independently assessed by three expert image analysts; for full details of this procedure see Rosen et al. (2017). Briefly, prior to rating images, all three raters were trained to >85% concordance with faculty consensus rating on an independent training sample of 100 images. Each rater evaluated every raw T1 image on a 0—2 Likert scale, where unusable images were coded as “0”, usable images with some artifact were coded as “1”, and images with none or almost no artifact were coded as “2”. All images with an average rating of 0 were excluded from analyses (*n* = 3); of the remaining images in the final sample *n* = 2 had an average manual rating of 0.67, *n* = 16 were rated as 1, *n* = 16 were rated as 1.33, *n* = 35 were rated as 1.67, and the remaining *n* = 358 had an average rating of 2. As described below, these average manual quality ratings were included in sensitivity analyses. In addition, we examined the distribution of cortical thickness values within anatomically defined regions created using a multi-atlas labeling technique (see below). For each region, we created a distribution of thickness values; subjects with a cortical thickness value >2 SD from the mean were flagged for that region. This procedure was repeated for all 98 cortical regions, and the number of flags was summed across regions; this summarized the number of regions per subject that had an outlying value. Subjects with >2.5 S.D. number of regional outliers were flagged for manual re-evaluation. Notably, this extensive post-processing QA procedure did initially identify 1 subject who failed the ANTs CT procedure. For this subject, minor parameter adjustments were made and the procedure was re-run, resulting in no subjects with major processing errors that required exclusion. Beyond this participant, there was no other manual intervention into standardized image processing procedures.

### Image processing and cortical thickness estimation

Structural image processing for estimating cortical thickness (CT) used tools included in Advanced Normalization Tools (ANTs, Tustison et al., 2014; RRID:SCR_004757). In order to avoid registration bias and maximize sensitivity to detect regional effects that can be impacted by registration error, a custom adolescent template and tissue priors were created. Structural images were then processed and registered to this template using the ANTs CT pipeline (Tustison et al., 2014). This procedure includes brain extraction, N4 bias field correction (Tustison et al., 2010), Atropos probabilistic tissue segmentation (Avants et al., 2011a), the top-performing SyN diffeomorphic registration method (Klein et al., 2010; Avants et al., 2011b; RRID:SCR_004757), and direct estimation of cortical thickness in volumetric space (Das et al., 2009). Large-scale evaluation studies have shown that this highly accurate procedure for estimating CT is more sensitive to individual differences over the lifespan than comparable techniques (Tustison et al., 2014). CT images were down-sampled to 2 mm voxels before applying non-negative matrix factorization, but no additional smoothing was performed.

### Non-negative matrix factorization

Cortical thickness was estimated as described above over the entire cortical surface. We sought to reduce CT in our sample into fewer dimensions, for two reasons. First, an efficient summary of CT data would allow us to evaluate only a small number of associations, rather than conduct voxel-wise inference that may be vulnerable to substantial Type I error (Eklund et al., 2016). Second, and importantly, prior work has shown that there are inherent patterns of covariance in CT (Zielinski et al., 2010; Alexander-Bloch et al., 2013; Sotiras et al., 2015; 2017), and analyzing the data according to this covariance structure may enhance interpretability.

Accordingly, we achieved both goals by using non-negative matrix factorization (NMF) to identify structural networks where cortical thickness co-varies consistently across individuals and brain regions (Sotiras et al., 2015). NMF has previously been shown to yield more interpretable and reproducible components than other decomposition techniques such as Principal Component Analysis or Independent Component Analysis (Sotiras et al., 2015; 2017). In contrast to the other techniques, NMF only yields compact networks with positive weights, which facilitates interpretation of effects.

The NMF algorithm takes as input a matrix *X* containing voxel-wise CT estimates (dimensions: 128,155 voxels × 427 participants), and approximates that matrix as a product of two matrices with non-negative elements: *X* ≅ *BC* (Figure 1). The first matrix, *B*, is of size *V* × *K* and contains the estimated non-negative networks and their respective loadings on each of the *V* voxels, where *K* is the user-specified number of networks. The *B* matrix (“CT loadings”) is composed of coefficients that denote the relative contribution of each voxel to a given network. These non-negative coefficients of the decomposition by necessity represent the entirety of the brain as a subject-specific addition of various parts. The second matrix, *C*, is of size *K* × *N* and contains subject-specific scores for each network. These subject-specific scores (“CT network scores”) indicate the contribution of each network in reconstructing the original CT map for each individual, and were evaluated for associations with DD as described below. We examined multiple NMF solutions ranging from 2 to 30 networks (in steps of 2) and calculated reconstruction error for each solution as the Frobenius norm between the CT data matrix and its NMF approximation (Sotiras et al., 2015; 2017). The optimal number of components was chosen based on the elbow of the gradient of the reconstruction error, such that the solution is adequate to model the structure of the data without modeling random noise (Sotiras et al., 2017). Network loadings were visualized on the inflated Population-Average, Landmark-, and Surface-based (PALS) cortical surfaces (Van Essen, 2005; RRID:SCR_002099) using Caret software (Van Essen et al., 2001; RRID:SCR_006260).

**Figure 1.**
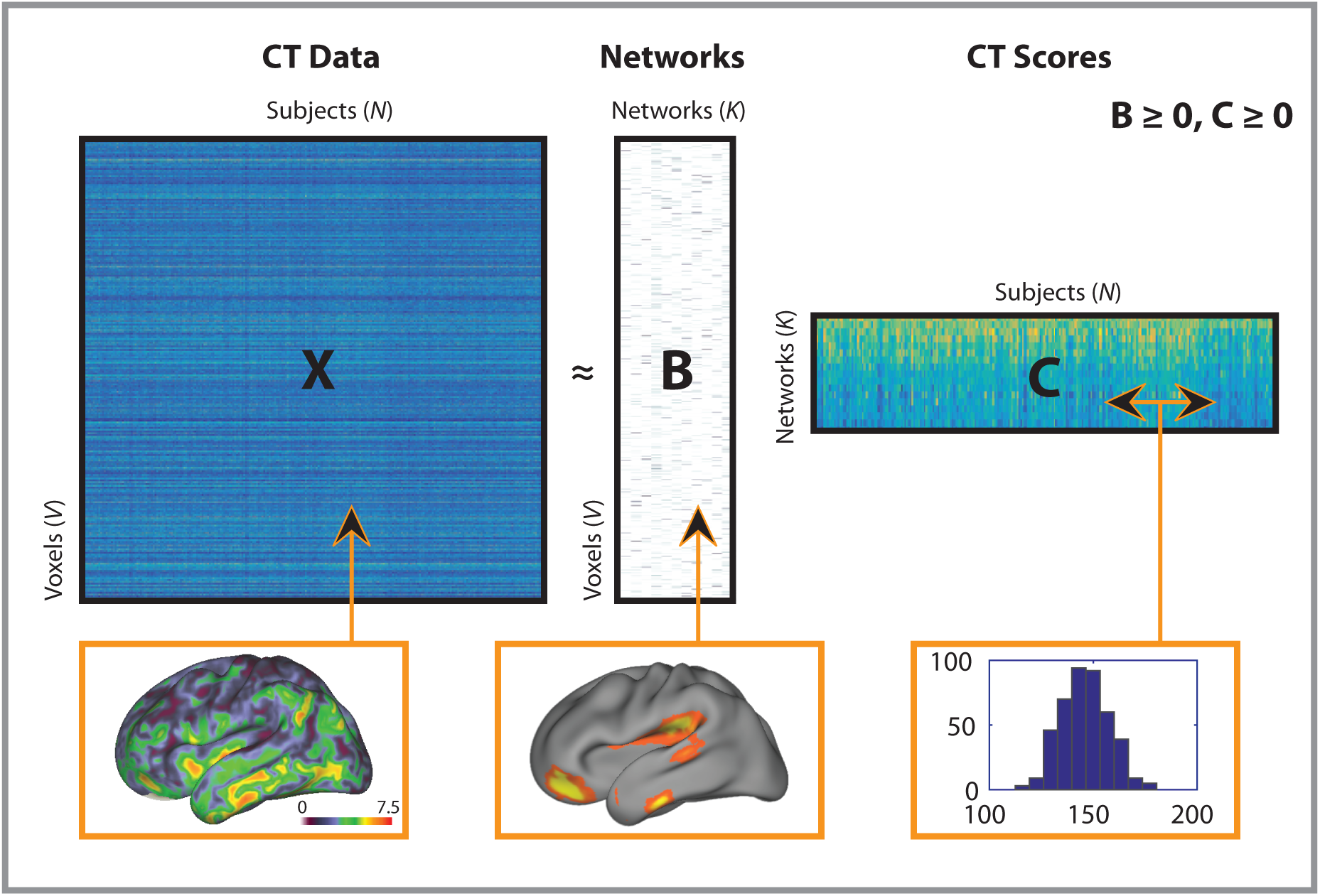
Schematic of non-negative matrix factorization and example data for each matrix. The *X* matrix represents the cortical thickness data (columns) for all subjects (rows); the CT map shows example CT data from one participant, and corresponds to a column in the *X* matrix. The *B* matrix represents estimated networks (columns) and their loadings on each voxel (rows); the example map shows loadings from one network, and corresponds to a column in the *B* matrix. The *C* matrix provides the subject-specific weights (columns) for each network (rows); the histogram shows CT scores in a single network, and corresponds to a row in the *C* matrix. Matrix sizes are shown with following dimensions: *V* = number of cortical thickness voxels, *N* = number of participants; *K* = number of networks.

### Regional parcellation using multi-atlas segmentation

In order to demonstrate that our results are robust to methodological variation, we also derived CT estimates in anatomically-defined regions of interest. We used a top-performing multi-atlas labelling approach to parcellate the brain into anatomical regions. This procedure has proven advantages over standard single-atlas approaches and has won open analysis challenges (Wang et al., 2012). Specifically, we used 24 young adult T1 images from the OASIS dataset (Marcus et al., 2007; RRID:SCR_007385), which have been manually labeled by Neuromorphometrics, Inc. (http://Neuromorphometrics.com/; RRID:SCR_005656). These images were each registered to each participant’s T1 image again using the top-performing SyN diffeomorphic registration method included in ANTs (Klein et al., 2010; Avants et al., 2011b; RRID:SCR_004757). Finally, a joint label fusion algorithm was used to synthesize the multiple warped atlas-labeled images into a final segmentation consisting of 98 gray matter regions (Wang et al., 2012). Mean thickness was calculated within each of these regions, and evaluated in group-level analyses identical to those conducted for NMF-derived networks, as described below.

### Experimental design and statistical analysis

To examine associations between DD and brain structure, we used a crosssectional sample of youths recruited as part of a large neurodevelopmental study. Brain development is frequently a nonlinear process (Giedd et al., 1999; Lenroot et al., 2007; Satterthwaite et al., 2014b). In order to capture both linear and nonlinear age effects, we modeled age with a penalized spline within Generalized Additive Models (GAMs; Wood, 2004; 2011; Vandekar et al., 2015). In this type of model, a penalty is assessed on nonlinearity using restricted maximum likelihood in order to avoid overfitting. GAMs were implemented in the R package ‘mgcv’ (https://cran.r-project.org/web/packages/mgcv/index.html; RRID:SCR_001905).

GAMs were first used to test for associations between DD and demographic variables such as age and sex. Next, we evaluated the association between DD and cognitive performance (as summarized by the overall cognitive performance factor and three domain-specific factor scores described above), while co-varying for sex and age. In both sets of analyses, DD was used as the dependent variable. Finally, univariate associations between DD and NMF-derived structural covariance networks were evaluated, with CT scores as the dependent variables and controlling for sex and age. Interactions between DD and age were evaluated but were not found to be significant, and were thus not included in the univariate models. To control multiple testing across either cognitive factors or structural covariance networks, we used the False Discovery Rate (FDR, Q<0.05; Benjamini and Hochberg, 1995).

In order to ensure that our results were not driven by potentially confounding factors, we conducted several sensitivity analyses. First, to ensure that our results were not driven by socio-economic status (SES), non-specific neurostructural effects, data quality or general cognitive abilities, we repeated these analyses while including maternal education, total brain volume, mean image quality rating, and the overall cognitive performance factor as model covariates in separate models. Second, we repeated our analyses while excluding participants who were taking a psychotropic medication at the time of scan (*n* = 52) or for whom medication data was not available (*n* = 3) to ensure that these participants did not bias the observed results.

### Multivariate analyses

The analyses described above examined univariate associations between each structural covariance network and DD. As a final step, we also investigated the multivariate predictive power of all cortical networks considered simultaneously, over and above that of two reduced models that included only demographics and non-neural correlates of DD (specifically, cognitive performance or maternal education). The first full model predicted DD using all 19 NMF networks, as well as age, sex, and the cognitive factors that were significantly associated with DD. The second full model predicted DD using all 19 NMF networks, as well as demographic variables including age, sex, and maternal education. These full models were compared to the reduced models (without the CT networks) using F-tests.

## RESULTS

### Impulsive choice is associated with reduced cognitive performance

Mean discount rate in our sample was 0.073 ± 0.088. Delay discounting was not related to demographic variables including age (*p* = 0.387). There was a non-significant trend toward more impulsive discounting in males (*p* = 0.07), and this trend was most prominent at younger ages (age by sex interaction: *p* = 0.09). In contrast, delay discounting was significantly associated with cognitive performance: youth who had higher discount rates also tended to have lower overall cognitive performance (partial *r* = −0.26, *p* < 0.0001). Follow-up analyses with a three-factor model describing specific cognitive domains revealed that this effect was driven primarily by an association with a combined executive functioning and complex reasoning factor (partial *r* = −0.29, p < 0.0001). Greater discounting was also associated with diminished memory accuracy (partial *r* = −0.20, p < 0.0001), whereas there was no significant relationship between DD and social cognition (partial *r* = −0.08, p = 0.10).

### Non-negative matrix factorization identifies structural covariance networks

Next, we sought to identify structural covariance networks in CT using NMF. NMF provides a data-driven way to identify structural covariance networks, where cortical thickness varies in a consistent way across individuals. As NMF identifies structural networks at a resolution set by the user, we examined solutions ranging from 2 to 30 networks (in steps of 2). As expected, reconstruction error consistently decreased as the number of networks increased. Similar to previous applications of this method (Sotiras et al., 2015), reconstruction error stabilized at 20 networks (**Figure 2**). Accordingly, the 20-network solution was used for all subsequent analyses (**Figure 3**).

**Figure 2.**
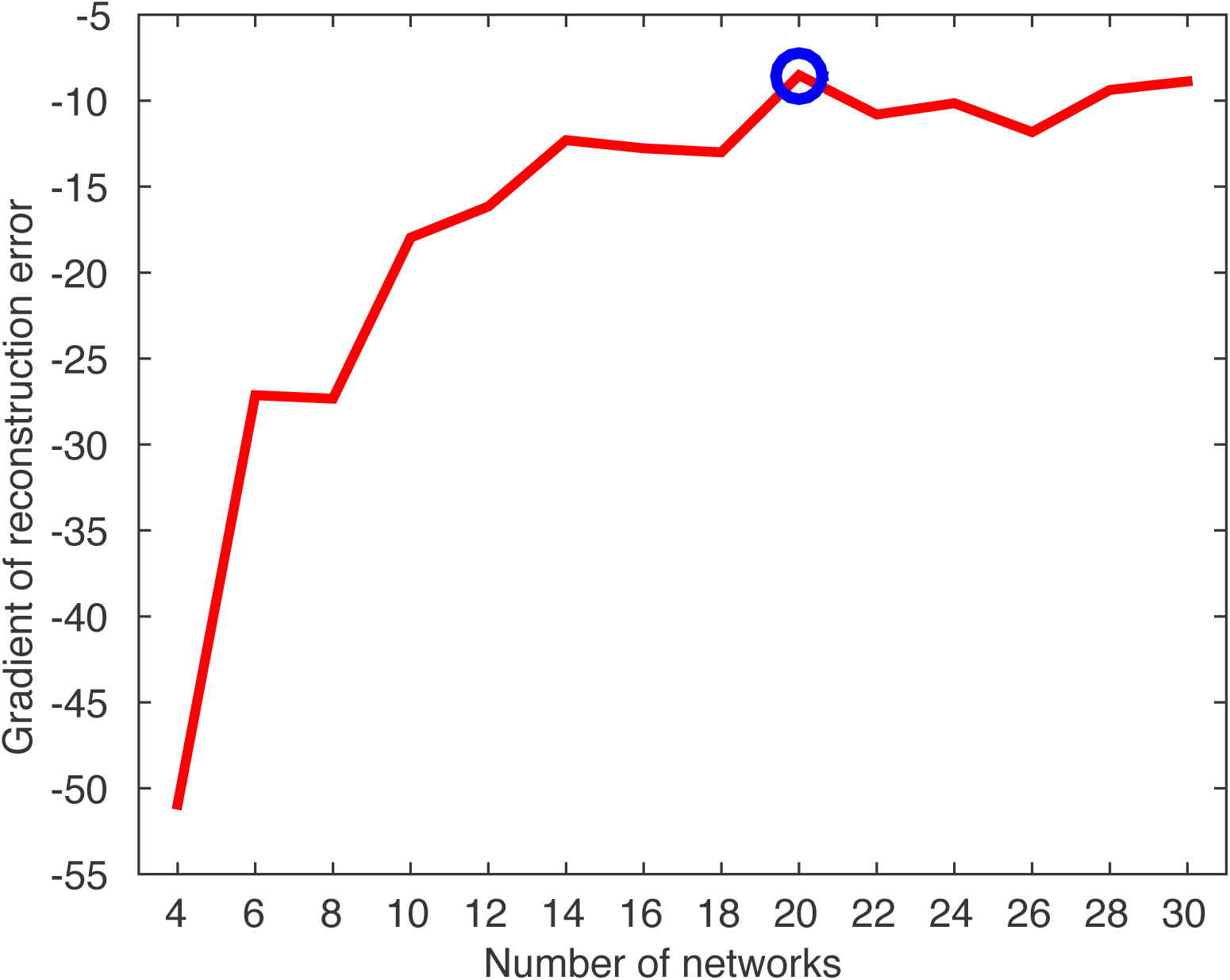
NMF reconstruction error identifies 20 cortical networks as the optimal resolution for cortical thickness data. Plot of reconstruction error gradient for NMF at multiple resolutions; the gradient is the difference in reconstruction error as the NMF solution increases by 2 networks. Blue circle indicates selected NMF solution of 20 networks.

**Figure 3.**
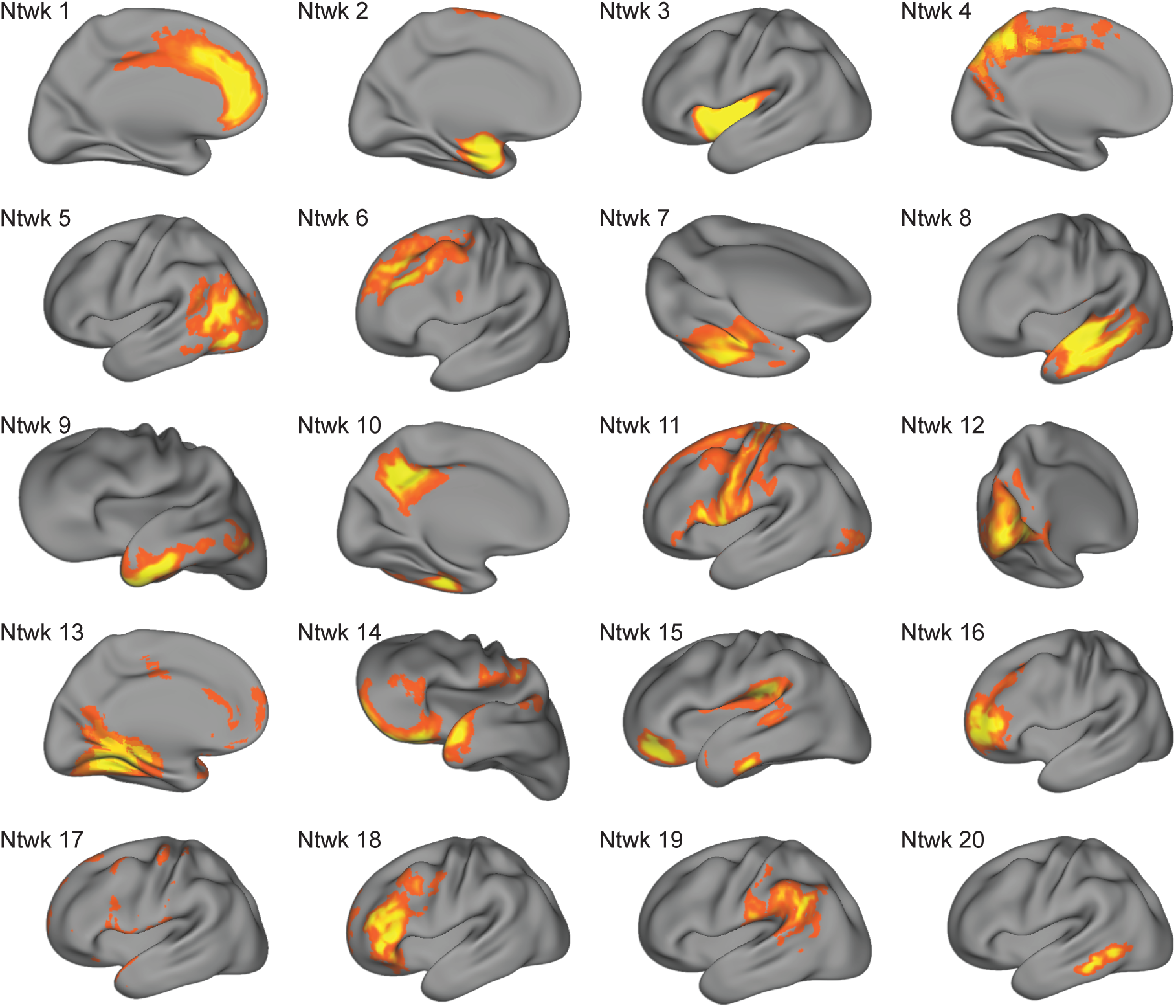
Structural covariance networks delineated by NMF. Visualization of structural covariance networks from the 20-network NMF solution. The spatial distribution of each network is indicated by loadings at each voxel in arbitrary units (from *B* matrix in NMF factorization); warmer colors represent higher loadings. For each network, we show one view that best captures the main area(s) of coverage. Approximate anatomical coverage of each structural covariance network: 1) medial prefrontal cortex and cingulate cortex; 2) medial temporal lobe; 3) insula; 4) medial posterior parietal cortex, including the precuneus; 5) temporo-occipital cortex; 6) dorsolateral prefrontal cortex (dlPFC); 7) fusiform gyrus; 8) lateral temporal lobe; 9) lateral temporal lobe and temporal pole; 10) posterior cingulate cortex and temporal lobe; 11) frontal and parietal cortex, including primary motor and somatosensory cortices; 12) occipital cortex; 13) medial temporal cortex, anterior cingulate cortex (ACC) and posterior cingulate cortex (PCC); 14) orbitofrontal cortex (OFC), frontal and temporal poles; 15) ventromedial prefrontal cortex (vmPFC), inferior temporal lobe, auditory cortex, temporoparietal junction (TPJ); 16) dorsal OFC; 17) the dura matter, a noise component that was not evaluated further; 18) dlPFC; 19) angular and supramarginal gyri; 20) posterior inferior temporal lobe.

As in prior work using NMF (Sotiras et al., 2017), the structural covariance networks identified were highly symmetric bilaterally. Networks included specific cortical regions that are relevant to reward processing and decision-making, such as ventromedial prefrontal cortex (vmPFC) and orbitofrontal cortex (OFC). Notably, when combined, several of the networks corresponded to aspects of functional brain networks. For example, networks 1 and 3 loaded on ACC and anterior insula, respectively, similar to the "salience network” (Seeley et al., 2007). Furthermore, specific networks defined lower-order systems, including motor (network 11) and visual (network 12) cortex. The 20-network solution also included a noise component (network 17), which was subsequently excluded from all analyses, resulting in 19 networks evaluated in total.

### Greater delay discounting is associated with diminished cortical thickness

Having identified 19 interpretable structural covariance networks using NMF, we next examined associations with DD while controlling for sex as well as linear and nonlinear age effects using penalized splines. Univariate analyses revealed that there was a significant association (after FDR correction) in eleven networks (**Table 1**). In each of these networks, impulsive choice, indicated by high discount rates, was associated with diminished cortical thickness. Notably, the strongest effects were found in two networks comprised of the ventromedial prefrontal cortex and orbitofrontal cortex, both regions known to be critical for reward-related decision-making. These two networks also included parts of the temporal pole and temporoparietal junction, TPJ (networks 14 and 15; **Figure 4**). Other networks where DD was associated with reduced CT included the temporal poles (network 9), lateral (network 8) and posterior temporal (network 20) lobes, dorsolateral prefrontal cortex (network 18), insula (network 3), fusiform gyrus (network 7), fronto-parietal cortex (network 11), and visual cortex (network 12).

**Figure 4.**
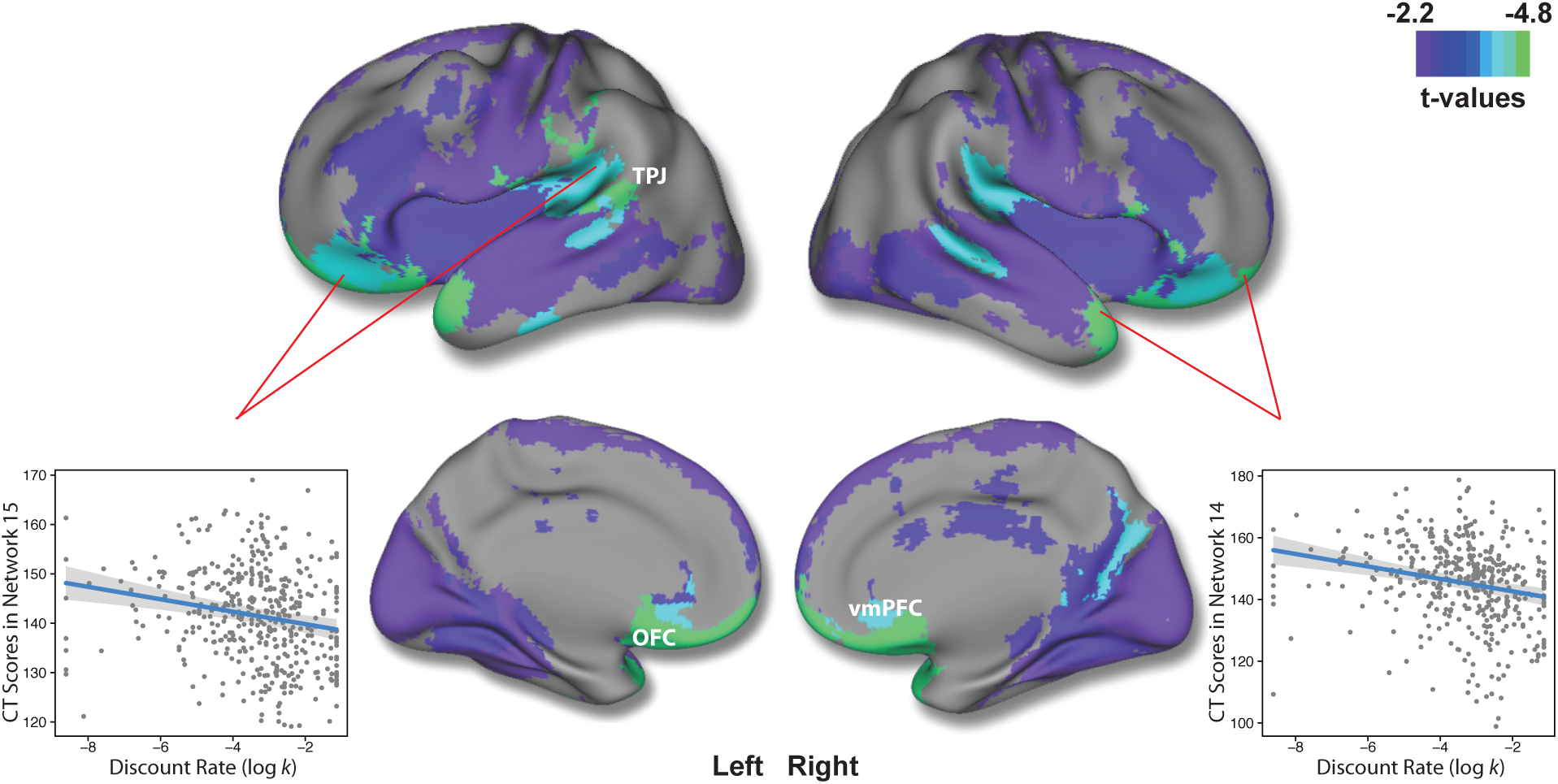
Higher discounting is associated with diminished cortical thickness in frontal, temporal, and parietal areas. Regions of FDR-significant association between log *k* and structural covariance networks. The composite network visualization was obtained by assigning each voxel to the network which has the highest loading for that voxel (from the *B* matrix), across all 19 networks. Maximal effects were observed in Networks 14 and 15, which included orbitofrontal cortex and ventromedial prefrontal cortex. Scatterplots for log *k*-CT association in these networks are shown, while adjusting for model covariates. Gray envelope represents the 95% confidence interval.

**Table 1.**
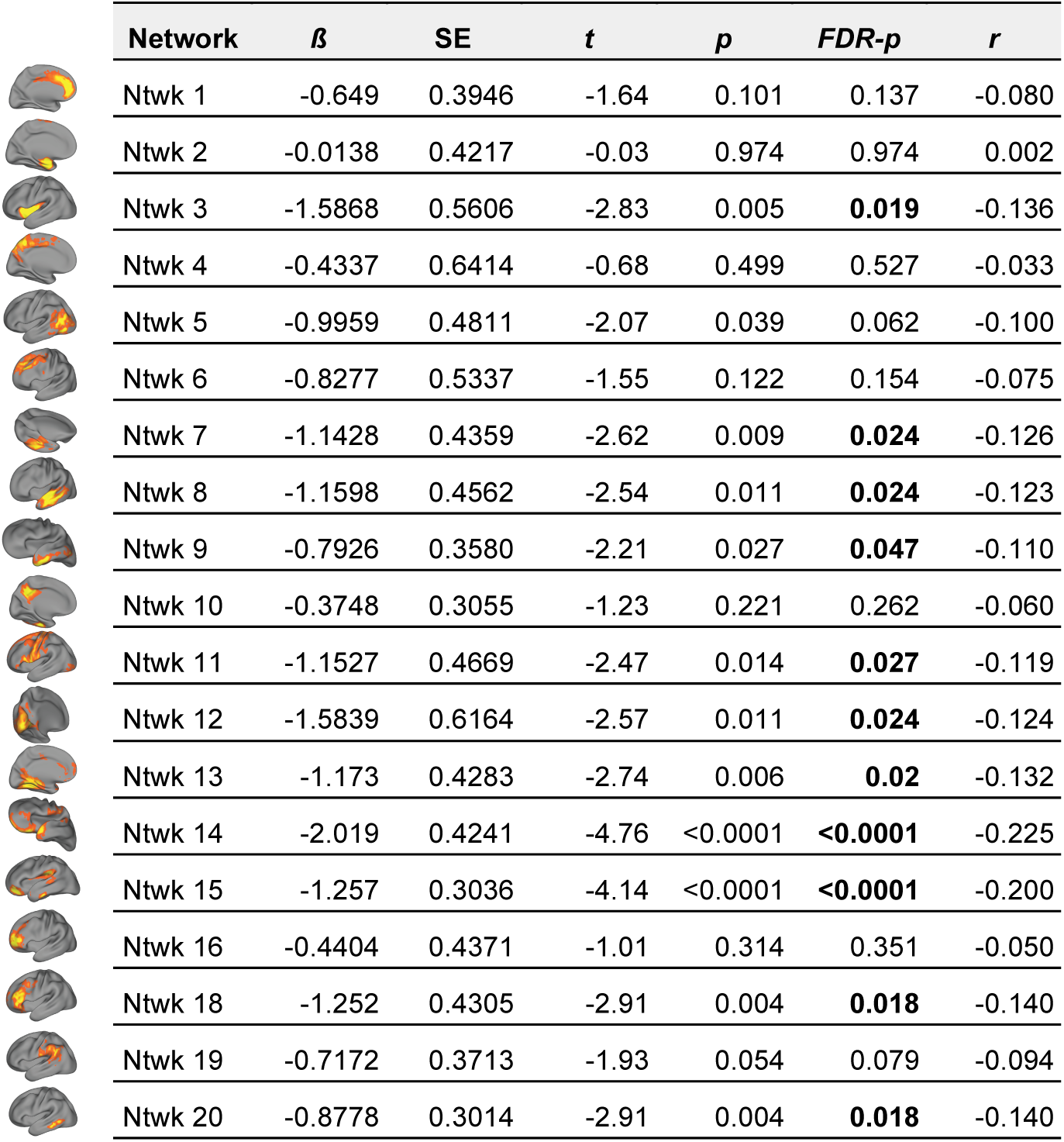
**Association between delay discounting and NMF-derived structural covariance networks.** β (unstandardized regression coefficient), SE (β’s standard error), t (t-value for testing β against 0, *dfs* = 423), p-value, and FDR-corrected p-values are obtained from separate general additive models run for each network. In this model, discount rate (log *k*) predicts cortical thickness scores, controlling for age (fit as a penalized spline) and sex. As an estimate of the linear effect size, *r* is the partial Pearson’s correlation coefficient between discount rate and CT scores in each network, while adjusting for linear age, quadratic age, and sex. FDR-significant p-values are indicated in bold.

| | Network | $\beta$ | SE | $t$ | $p$ | FDR-$p$ | $r$ |
| --- | --- | --- | --- | --- | --- | --- | --- |
|  | Ntwk 1 | -0.649 | 0.3946 | -1.64 | 0.101 | 0.137 | -0.080 |
|  | Ntwk 2 | -0.0138 | 0.4217 | -0.03 | 0.974 | 0.974 | 0.002 |
|  | Ntwk 3 | -1.5868 | 0.5606 | -2.83 | 0.005 | **0.019** | -0.136 |
|  | Ntwk 4 | -0.4337 | 0.6414 | -0.68 | 0.499 | 0.527 | -0.033 |
|  | Ntwk 5 | -0.9959 | 0.4811 | -2.07 | 0.039 | 0.062 | -0.100 |
|  | Ntwk 6 | -0.8277 | 0.5337 | -1.55 | 0.122 | 0.154 | -0.075 |
|  | Ntwk 7 | -1.1428 | 0.4359 | -2.62 | 0.009 | **0.024** | -0.126 |
|  | Ntwk 8 | -1.1598 | 0.4562 | -2.54 | 0.011 | **0.024** | -0.123 |
|  | Ntwk 9 | -0.7926 | 0.3580 | -2.21 | 0.027 | **0.047** | -0.110 |
|  | Ntwk 10 | -0.3748 | 0.3055 | -1.23 | 0.221 | 0.262 | -0.060 |
|  | Ntwk 11 | -1.1527 | 0.4669 | -2.47 | 0.014 | **0.027** | -0.119 |
|  | Ntwk 12 | -1.5839 | 0.6164 | -2.57 | 0.011 | **0.024** | -0.124 |
|  | Ntwk 13 | -1.173 | 0.4283 | -2.74 | 0.006 | **0.02** | -0.132 |
|  | Ntwk 14 | -2.019 | 0.4241 | -4.76 | <0.0001 | **<0.0001** | -0.225 |
|  | Ntwk 15 | -1.257 | 0.3036 | -4.14 | <0.0001 | **<0.0001** | -0.200 |
|  | Ntwk 16 | -0.4404 | 0.4371 | -1.01 | 0.314 | 0.351 | -0.050 |
|  | Ntwk 18 | -1.252 | 0.4305 | -2.91 | 0.004 | **0.018** | -0.140 |
|  | Ntwk 19 | -0.7172 | 0.3713 | -1.93 | 0.054 | 0.079 | -0.094 |
|  | Ntwk 20 | -0.8778 | 0.3014 | -2.91 | 0.004 | **0.018** | -0.140 |

### Association between cortical thickness and delay discounting is independent of age-related changes in cortical thickness

Having established that individual differences in DD are associated with CT, we next examined whether this effect was moderated by age. Notably, there was no significant age by DD interaction in any network (median *p* = 0.77, range: 0.09—0.94). Thus, age-related changes in CT were similar in both high and low discounters, but those with higher discount rates had thinner cortex across the age range examined (**Figure 5**).

**Figure 5.**
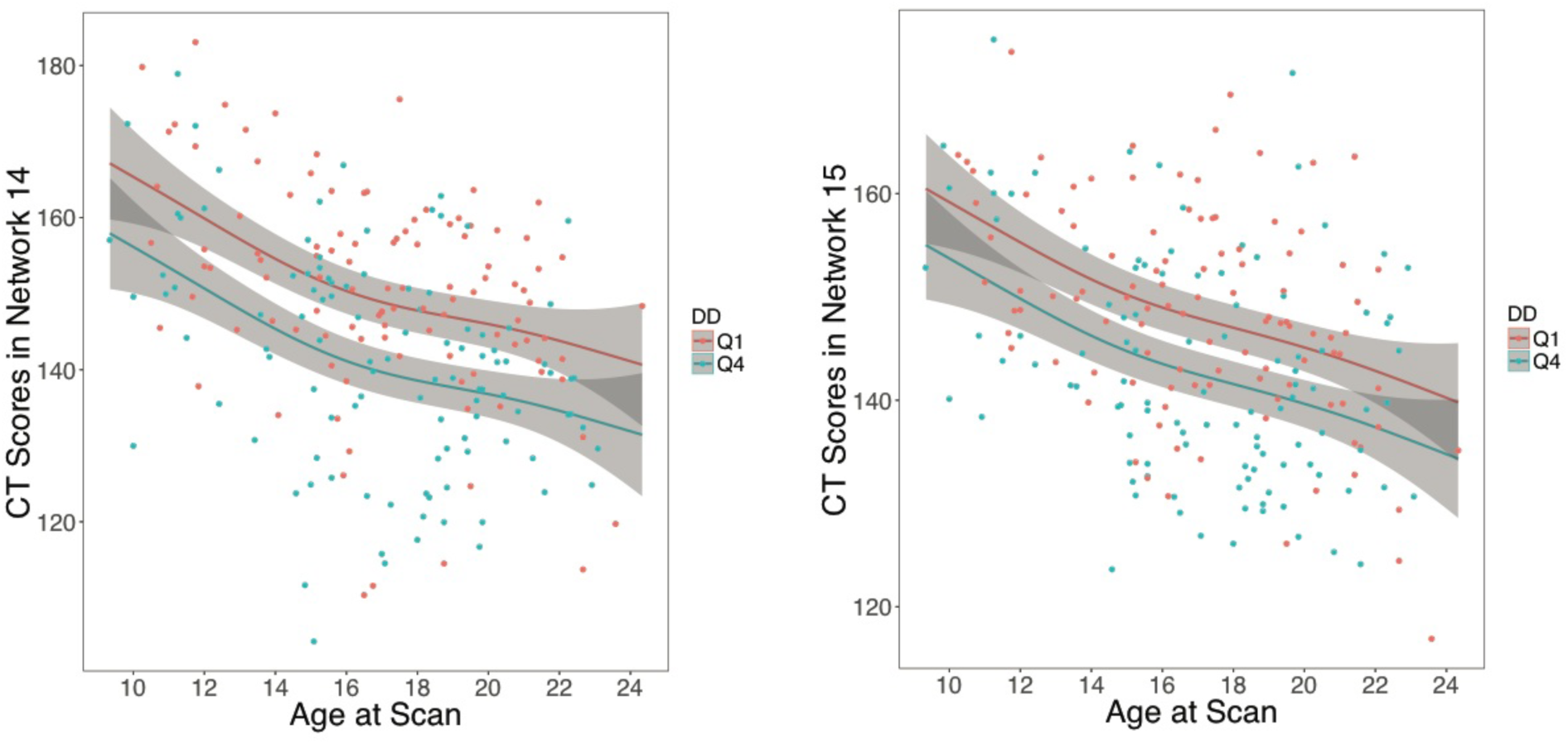
Association between cortical thickness and delay discounting is independent of age. Scatterplots for relationship between age and CT in networks 14 and 15, separated by top (Q4) and bottom (Q1) quartiles of log *k*. The Q4 quartile group contains participants with the most impulsive preferences. For each quartile, the age-CT relationship is shown after adjusting for model covariates, and includes the 95% confidence intervals (gray envelopes).

### Sensitivity analyses provide convergent results

We conducted sensitivity analyses to evaluate potentially confounding variables including maternal education, total brain volume, image data quality, general cognitive abilities, and psychotropic medications. First, we examined if the results could be explained by differences in maternal education, a proxy of socioeconomic status. Discount rate was significantly associated with maternal education (partial *r* = −0.164, *p* = 0.0007), but including it in the model did not have a great impact on results.

Specifically, 7 of 11 networks found to be related to DD remained FDR-significant, including the vmPFC and OFC networks; the other 4 networks trended towards significance (*p_fdr_* < 0.067). Second, we examined the effect of total brain volume on our findings. After adding total brain volume as a covariate, 10 of 11 networks remained FDR-significant for association with DD, with the remaining network showing a trend towards FDR-significance (*p_fdr_* = 0.0762). Third, we included mean image quality rating (averaged across three expert raters) as a model covariate. Despite the fact that data quality was significantly related to discount rate (Spearman’s partial *rho* = −0.159, *p* = 0.001), 7 of 11 networks continued to have an FDR-significant association after inclusion of this covariate (including the vmPFC and OFC networks), and 3 out of 11 networks had FDR-corrected p-values of < 0.10. Fourth, we examined the effect of cognitive abilities, as measured by the overall cognitive performance factor. After including this variable as a covariate, 5 of 11 networks related to DD remained FDR-significant, including networks spanning the vmPFC, OFC, insula, and inferior temporal lobe. Finally, we repeated this analysis after excluding 52 participants who were taking psychotropic medication at the time of scan and 3 participants for whom medication data were missing. Despite the reduced power of this smaller sample, 10 of 11 networks remained FDR-significant, with the final network showing a trend towards significance (*p_fdr_* = 0.0503).

### Analyses with anatomically defined regions yield convergent results

To evaluate the robustness of the relationship between DD and CT to methodological variation, we also examined associations within 98 anatomically-based regions. Univariate analyses controlling for sex and age revealed significant negative associations between DD and CT in 24 of these regions (**Figure 6**). Consistent with the previously described NMF results, impulsive choice— indexed by greater discounting— was associated with diminished cortical thickness in medial frontal cortex, orbitofrontal cortex, fusiform gyrus, frontal and temporal poles, insular cortex, middle and superior temporal gyri, precentral gyrus, and occipital cortex.

**Figure 6.**
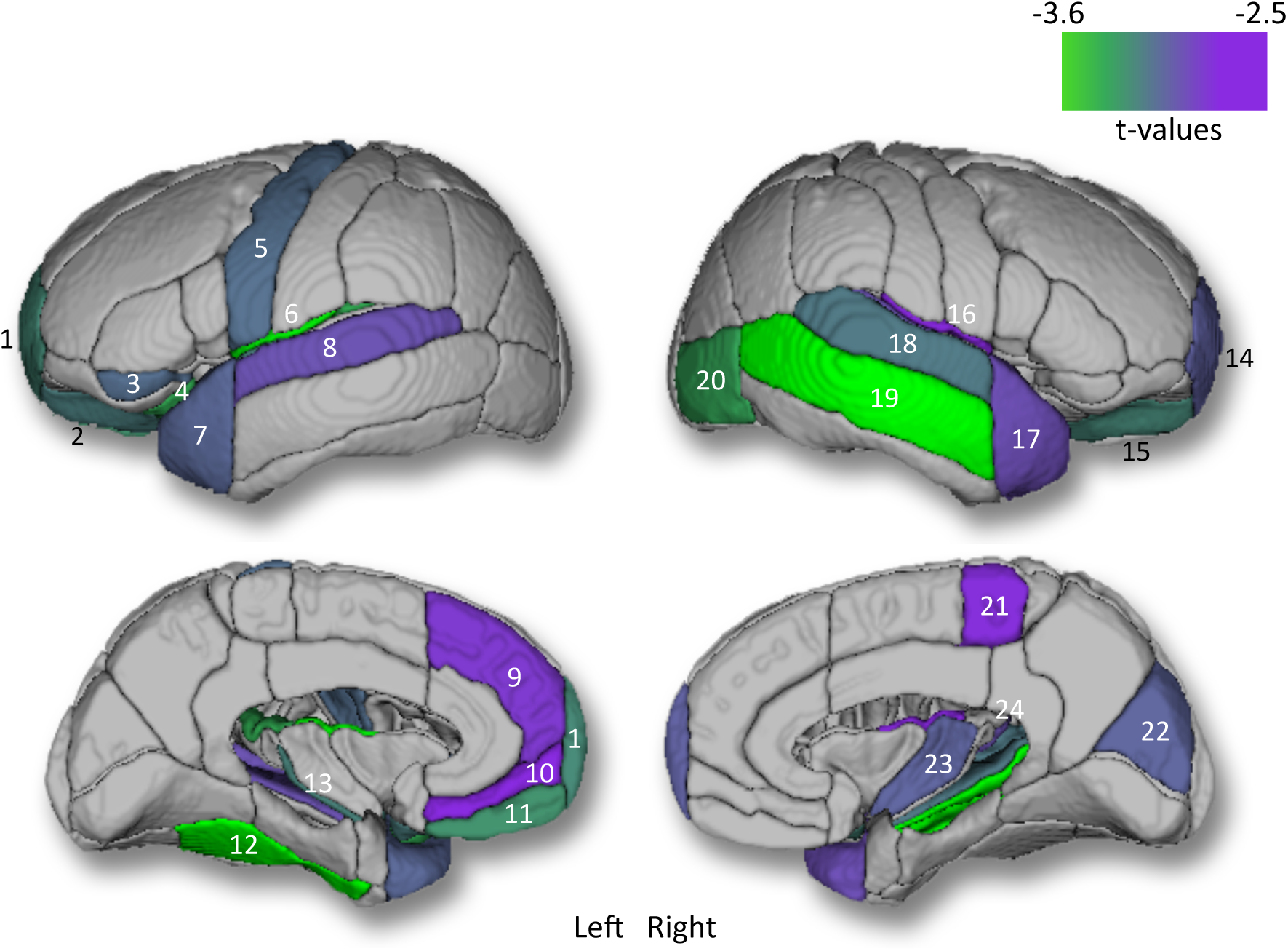
Higher discounting is associated with diminished regional cortical thickness in frontal and temporal regions. FDR-significant associations between log *k* and CT estimated in anatomically-defined regions. Significant regions include the following: left frontal pole (1); left medial orbital gyrus (2); left orbital part of the inferior frontal gyrus (3); left posterior orbital gyrus (4); left precentral gyrus (5); left central operculum (6); left temporal pole (7); left superior temporal gyrus (8); left superior frontal gyrus, medial segment (9); left medial frontal cortex (10); left gyrus rectus (11); left fusiform gyrus (12); left planum polare (13); right frontal pole (14); right medial orbital gyrus (15); right central operculum (16); right temporal pole (17); right superior temporal gyrus (18); middle temporal gyrus (19); right inferior occipital gyrus (20); right precentral gyrus, medial segment (21); right cuneus (22); right posterior insula (23); right planum temporale (24).

### Covariance networks provide improved prediction of DD over demographic and cognitive data

The univariate analyses described above demonstrated that reduced CT in several structural covariance networks is associated with impulsive choice. Next, we tested whether a multivariate model including all structural networks could accurately predict DD on an individual basis. Delay discounting predicted from a model of CT scores in all 19 networks, as well demographic data (age and sex), was significantly correlated with actual delay discounting behavior (*r* = 0.33, *p* < 0.0001; **Figure 7**). Adding CT scores to a reduced model with demographics alone improved model fit (*F(_405,424_)* = 2.37, *p* = 0.001); DD predicted from this reduced model with demographics only achieved a correlation of 0.097 (*p* = 0.043) with actual log *k* values.

**Figure 7.**
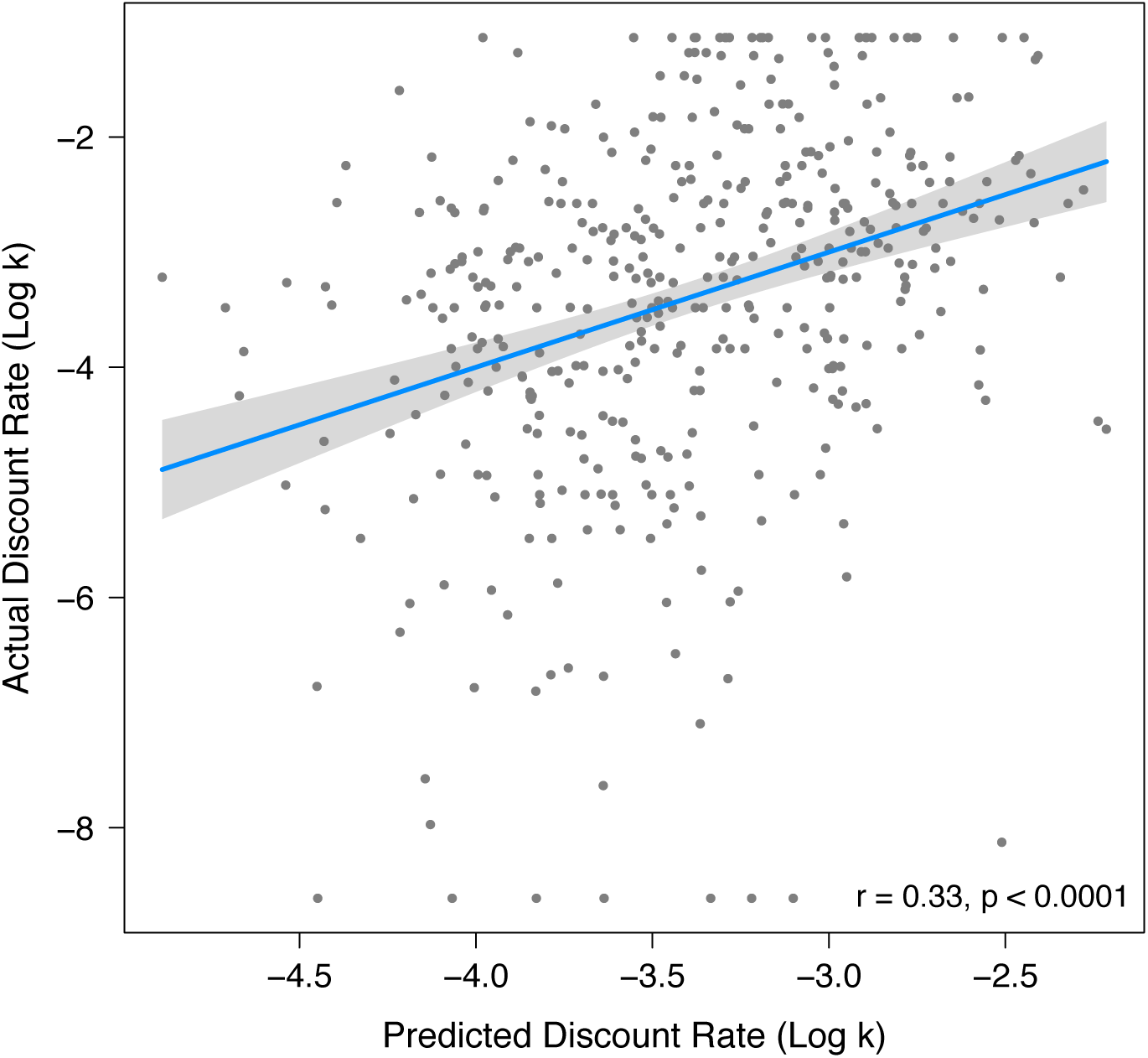
CT data from structural covariance networks predicts delay discounting. Scatterplot for relationship between actual log *k* values and predicted log *k* from multivariate CT prediction. Multivariate prediction is based on CT scores from all structural covariance networks plus demographic variables: sex and age. Scatterplots include line of best fit for this association with a 95% confidence interval (gray envelope).

Importantly, CT data also improved prediction above and beyond that achieved by cognitive predictors: adding CT scores to a model with cognitive performance as well as demographics improved the model fit (*F(_403,422_)* = 1.63, *p* = 0.047). DD predicted from the reduced model with just demographics and cognition achieved a correlation of 0.31 (*p* < 0.0001) between model-predicted and actual log *k* values, compared to a correlation of 0.40 (*p* < 0.0001) from a complete model including CT data, cognitive data, and demographics. Furthermore, CT networks also improved prediction above and beyond that achieved by maternal education, a proxy of socioeconomic status, which was correlated with DD (*F*(_401,420_) = 1.97, *p* = 0.009). DD predicted from a model with demographic variables such as age, sex, and maternal education achieved a correlation of 0.19 (*p* < 0.0001) between predicted and actual *k* values, while *k* values estimated from the full model with CT data, demographics, and maternal education achieved a correlation of 0.34 (*p* < 0.0001) with actual *k* values.

## DISCUSSION

We examined associations between delay discounting and cortical thickness networks in a large adolescent sample. More impulsive preferences, as indexed by higher discounting, were associated with diminished CT in multiple networks. The strongest effects were found in OFC, vmPFC, temporal pole, and the TPJ. Associations between DD and brain structure did not vary over the age range studied, and could not be explained by confounding variables. Furthermore, consideration of structural networks improved prediction of DD above and beyond demographic and cognitive variables.

### Structural covariance networks related to DD overlap with known functional networks

Greater discounting was associated with decreased cortical thickness in multiple structural networks. Relative to previous reports of both neurofunctional and neurostructural correlates of DD (Peters and Büchel, 2011; Bernhardt et al., 2014; Kable and Levy, 2015), the effects we observed were fairly widespread across the brain. Notably, many of the regions encompassed by these networks correspond to findings from previous studies in adults, including functional networks known to be involved in DD. As hypothesized, we found associations between DD and CT in central elements of the valuation network, namely vmPFC (Bartra et al., 2013), the cognitive control network, including dlPFC (Peters and Büchel, 2011; Stanger et al., 2013), and the prospection network, involving medial temporal cortex (Peters and Büchel, 2011). While DD and CT relationships have not previously been evaluated in adolescents, one prior study documented diminished thickness in the ACC and medial PFC in association with greater DD in adults (Bernhardt et al., 2014). In addition to hypothesized effects, we also found associations between DD and CT in motor, somatosensory, and both early and higher-order visual cortices. Notably, when these effects were evaluated jointly in a multivariate model, CT networks enhanced prediction of DD above and beyond demographic and cognitive variables. This result contributes to efforts in neuroeconomics to improve prediction of decision-making behavior using brain-based measures obtained independently of the behavior itself (Kable and Levy, 2015), and suggests that structural covariance networks may be a useful marker of impulsive choice in youth.

### Results converge with data from lesion and neuromodulation studies

Although the negative associations between DD and CT were widespread and distributed, two structural covariance networks exhibited particularly strong associations with DD. Brain regions comprising these networks included vmPFC, OFC, temporal pole, and the TPJ. As mentioned above, our findings in vmPFC were expected based on substantial evidence from fMRI studies that this region is implicated in DD (Kable and Glimcher, 2007; Ballard and Knutson, 2009; Bartra et al., 2013). Furthermore, activity in vmPFC when merely thinking about the future predicts DD, such that lower discounters show greater activity when thinking about the far future (Cooper et al., 2013). Finally, consistent with our results, a study in adults reported that diminished CT in that region was associated with higher DD (Bernhardt et al., 2014).

Beyond the vmPFC, evidence suggests that regions including the OFC, temporal pole, and TPJ are both involved in and necessary for evaluating future outcomes in DD. First, lesion studies in patients with medial OFC damage show greater discounting of both primary and secondary rewards, compared to healthy controls and non-frontal damage patients (Sellitto et al., 2010), and this is the only region where injuries have been reported to increase discounting in humans. Notably, this relationship is dose-dependent, such that larger frontal lesions are associated with steeper discounting. Second, patients with semantic dementia, a disorder characterized by anterior temporal lobe atrophy, show greater discounting than controls (Chiong et al., 2015). Third, while the TPJ has typically been implicated in social cognition and theory of mind, recent data suggests that it also plays a role in both monetary and social discounting (Strombach et al., 2015; Soutschek et al., 2016). Importantly, disrupting the TPJ in healthy adults using transcranial magnetic stimulation increases discounting (Soutschek et al., 2016). Collectively, this evidence suggests that the disruption of OFC, anterior temporal lobe, and TPJ may promote impulsive choice.

### Associations with delay discounting are independent of age-related changes

While we replicated prior findings of association between lower discounting and higher IQ (Shamosh and Gray, 2008) and working memory (Shamosh et al., 2008), we did not find significant associations between DD and age (Scheres et al., 2006; Steinberg et al., 2009). This may be due to differences in sample composition, including relatively less dense sampling of younger ages and use of a dimensional rather than a stratified design that compared older and younger age groups. However, the lack of observed age effect is consistent with a recent review noting that age effects on DD are inconsistent and of a relatively small effect size (Romer et al., 2017). Notably, the association between brain structure and DD was stable across the entire age range surveyed in our sample. This result is consistent with a prior study of DD in adolescents and white matter integrity assessed using diffusion imaging (Olson et al., 2009). Together, these results imply that individual differences in brain structure associated with impulsive choice do not emerge specifically during adolescence. These results may also suggest that such individual differences in brain structure may emerge early in development, consistent with literature describing the importance of structural brain development in utero, during the peri-natal period, and during early childhood (Thomason et al., 2013; Di Martino et al., 2014). While speculative, future research may reveal that individual differences in brain structure which emerge early in life may impact evolving patterns of value and cognitive control system function in adolescence which, in turn, may contribute to impulsivity during this critical period (Casey et al., 2008; Bjork et al., 2010).

### Advantages of evaluating structural covariance networks in a large sample

The greater spatial extent of observed significant associations between brain structure and DD compared to prior results may be due to several aspects of our study. First, the large sample size afforded greater statistical power, and thus greater sensitivity, to detect effects in multiple networks. While the effect sizes of these associations were small, research documenting inflation of effect sizes in small studies suggests that our results are more likely to be an accurate reflection of the true effect size than data from modest samples (Button et al., 2013). Second, structural covariance networks defined by NMF provided a parsimonious summary of the high-dimensional imaging data. In contrast to anatomic atlases based on sulcal folding patterns, NMF identifies structural networks based on patterns of covariance in the data itself. This concise summary of the data limited multiple comparisons: we only evaluated 19 networks in our analyses, in contrast to the hundreds of thousands of voxels typically surveyed in mass-univariate voxel-based morphometry studies. This allowed us to use a rigorous FDR correction for all comparisons, rather than cluster-based inference that may produce substantial Type I error rates in many common implementations (Eklund et al., 2016).

### Limitations

Certain limitations of this study should be noted. First, the observed effects were independent of age, suggesting that differences in brain structure associated with impulsive choice may emerge earlier than the examined age range. Future investigations should consider longitudinal designs including early childhood to precisely capture the emergence of these effects. Second, we used hypothetical instead of real rewards in the DD task. However, prior studies have yielded similar results in both behavioral (Johnson and Bickel, 2002) and functional neuroimaging paradigms (Bickel et al., 2009). Third, we cannot completely rule out potential confounding variables which may be correlated with DD. Previous studies have described associations between CT and SES in adolescence (Mackey et al., 2015), though importantly our results remained largely unaffected after controlling for maternal education, a proxy of SES. Fourth, while the ANTs DiReCT method of quantifying CT has been shown as highly accurate and more discriminative than comparable techniques in large-scale evaluation studies (Tustison et al., 2014), it does not allow high-resolution voxel-wise analyses or provide information (such as surface area) regarding potentially important subcortical structures.

### Conclusions and future directions

Understanding impulsive choice in adolescence is important because impulsivity is associated with a host of risky behaviors and outcomes, such as tobacco use (Reynolds, 2004), alcohol use (Fernie et al., 2013), obesity (Fields et al., 2013), and early sexual initiation (Khurana et al., 2012), which lead to substantial morbidity and mortality during adolescence. Leveraging a large developmental sample and advanced analytics, we found that individual variability in brain structure explains differences in DD in adolescence. Taken together, our results indicate that higher DD in youth is associated with reduced cortical thickness in multiple networks, including those known to be essential for valuation. These results emphasize that risky behaviors in adolescents should be considered in the context of individual differences of structural brain networks that are present early in life. Moving forward, such brain-based measures could potentially be used as biomarkers to identify youth at particularly high risk for negative outcomes. Future studies should evaluate associations between DD, brain structure, and psychopathology. Such efforts could potentially aid in stratifying youth within targeted clinical trials aiming to reduce impulsivity and risk-taking behaviors during this critical period.

## ACKNOWLEDGEMENTS

We thank the acquisition and recruitment team, including Karthik Prabhakaran and Jeff Valdez. Thanks to Chad Jackson for data management and systems support and Monica Calkins for phenotyping expertise. Supported by grants from the National Institute of Mental Health: R01MH107703 (TDS). The PNC was funded through NIMH RC2 grants MH089983 and MH089924 (REG). Additional support was provided by MH098899 and DA029149 (JWK), K01MH102609 (DRR), R01MH101111 (DHW), R01-AG014971 and RF1- AG054409 (CD & AS), P50MH096891 (REG), R01NS085211 (RTS), R01MH107235 (RCG), the Dowshen Program for Neuroscience, and the Lifespan Brain Institute at the Children’s Hospital of Philadelphia and Penn Medicine. Seed grant support by the Center for Biomedical Computing and Image Analysis (CBICA) was awarded for developing statistical analyses (RTS & TDS), as well as non-negative matrix factorization software (AS & TDS). We used images from the OASIS project, which was supported by the following grants: P50 AG05681, P01 AG03991, R01 AG021910, P20 MH071616, U24 RR021382.

## REFERENCES

Alexander-Bloch A, Giedd JN, Bullmore E (2013) Imaging structural co-variance between human brain regions. Nat Rev Neurosci 14:322–336.

Audrain-McGovern J, Rodriguez D, Epstein LH, Cuevas J, Rodgers K, Wileyto EP (2009) Does delay discounting play an etiological role in smoking or is it a consequence of smoking?. Drug Alcohol Depend 103:99–106.

Avants BB, Tustison NJ, Wu J, Cook PA, Gee JC (2011a) An open source multivariate framework for n-tissue segmentation with evaluation on public data. Neuroinformatics 9:381–400.

Avants BB, Tustison NJ, Song G, Cook PA, Klein A, Gee JC (2011b) A reproducible evaluation of ANTs similarity metric performance in brain image registration. Neuroimage 54:2033–2044.

Ballard K, Knutson B (2009) Dissociable neural representations of future reward magnitude and delay during temporal discounting. Neuroimage 45:143–150.

Bartra O, McGuire JT, Kable JW (2013) The valuation system: a coordinate-based meta-analysis of BOLD fMRI experiments examining neural correlates of subjective value. Neuroimage 76:412–427.

Benjamini Y, Hochberg Y (1995) Controlling the false discovery rate: a practical and powerful approach to multiple testing. Journal of the Royal Statistical Society. Series B (Methodological), 289–300.

Bernhardt BC, Smallwood J, Tusche A, Ruby FJ, Engen HG, Steinbeis N, Singer T (2014) Medial prefrontal and anterior cingulate cortical thickness predicts shared individual differences in self-generated thought and temporal discounting. Neuroimage 90:290–297.

Bickel WK, Miller ML, Yi R, Kowal BP, Lindquist DM, Pitcock JA (2007) Behavioral and neuroeconomics of drug addiction: competing neural systems and temporal discounting processes. Drug Alcohol Depend 90:S85–S91.

Bickel WK, Pitcock JA, Yi R, Angtuaco EJ (2009) Congruence of BOLD response across intertemporal choice conditions: fictive and real money gains and losses. J Neurosci 29:8839–8846.

Bjork JM, Momenan R, Hommer DW (2009) Delay discounting correlates with proportional lateral frontal cortex volumes. Biol Psychiatry 65:710–713.

Bjork JM, Smith AR, Chen G, Hommer DW (2010) Adolescents, adults and rewards: comparing motivational neurocircuitry recruitment using fMRI. PloS one, 5(7), e11440.

Button KS, Ioannidis JP, Mokrysz C, Nosek BA, Flint J, Robinson ES, Munafò MR (2013) Power failure: why small sample size undermines the reliability of neuroscience. Nat Rev Neurosci 14:365–376.

Casey BJ, Jones RM, Hare TA (2008) The adolescent brain. Ann N Y Acad Sci 1124:111–126.

Chiong W, Wood KA, Beagle AJ, Hsu M, Kayser AS, Miller BL, Kramer JH (2015) Neuroeconomic dissociation of semantic dementia and behavioural variant frontotemporal dementia. Brain 139:578–587.

Cho SS, Pellecchia G, Aminian K, Ray N, Segura B, Obeso I, Strafella AP (2013) Morphometric correlation of impulsivity in medial prefrontal cortex. Brain Topogr 26:479–487.

Cooper N, Kable JW, Kim BK, Zauberman G (2013) Brain activity in valuation regions while thinking about the future predicts individual discount rates. J Neurosci 33:13150–13156.

Das SR, Avants BB, Grossman M, Gee JC (2009) Registration based cortical thickness measurement. Neuroimage 45:867–879.

Di Martino A, Fair DA, Kelly C, Satterthwaite TD, Castellanos FX, Thomason ME, Craddock RC, Luna B, Leventhal BL, Zuo XN, Milham MP (2014) Unraveling the miswired connectome: a developmental perspective. Neuron 83:1335–1353.

Dombrovski AY, Siegle GJ, Szanto K, Clark L, Reynolds CF, Aizenstein H (2012) The temptation of suicide: striatal gray matter, discounting of delayed rewards, and suicide attempts in late-life depression. Psychol Med 42:1203–1215.

Eaton DK, Kann L, Kinchen S, Shanklin S, Flint KH, Hawkins J, Harris WA, Lowry R, McManus T, Chyen D, Whittle L, Lim C, Wechsler H (2012) Youth risk behavior surveillance-United States, 2011. Morbidity and Mortality Weekly Report. Surveillance Summaries (Washington, DC: 2002) 61:1–162.

Eklund A, Nichols TE, Knutsson H (2016) Cluster failure: why fMRI inferences for spatial extent have inflated false-positive rates. Proc Natl Acad Sci U S A 113:7900–7905.

Fernie G, Peeters M, Gullo MJ, Christiansen P, Cole JC, Sumnall H, Field M (2013) Multiple behavioural impulsivity tasks predict prospective alcohol involvement in adolescents. Addiction 108:1916–1923.

Field M, Christiansen P, Cole J, Goudie A (2007) Delay discounting and the alcohol Stroop in heavy drinking adolescents. Addiction 102:579–586.

Fields SA, Sabet M, Reynolds B (2013) Dimensions of impulsive behavior in obese, overweight, and healthy-weight adolescents. Appetite 70:60–66.

Giedd JN, Blumenthal J, Jeffries NO, Castellanos FX, Liu H, Zijdenbos A, Paus T, Evans, AC, Rapoport JL (1999) Brain development during childhood and adolescence: a longitudinal MRI study. Nat Neurosci 2:861–863.

Gur RC, Richard J, Hughett P, Calkins ME, Macy L, Bilker WB, Brensinger C, Gur RE (2010) A cognitive neuroscience-based computerized battery for efficient measurement of individual differences: standardization and initial construct validation. J Neurosci Meth 187:254–262.

Gur RC, Richard J, Calkins ME, Chiavacci R, Hansen JA, Bilker WB, Loughead J, Connolly JJ, Qiu H, Mentch FD, Abou-Sleiman PM, Hakonarson H, Gur RE (2012) Age group and sex differences in performance on a computerized neurocognitive battery in children age 8– 21. Neuropsychology 26:251–265.

Johnson MW, Bickel WK (2002) Within-subject comparison of real and hypothetical money rewards in delay discounting. J Exp Anal Behav 77:129–146.

Kable JW (2013) Valuation, Intertemporal Choice, and Self-Control. In: Neuroeconomics: Decision making and the brain (Glimcher PW, Fehr E, ed), pp173–192. Elsevier Inc.

Kable JW, Glimcher PW (2007) The neural correlates of subjective value during intertemporal choice. Nat Neurosci 10:1625–1633.

Kable JW, Levy I (2015) Neural markers of individual differences in decision-making. Curr Opin Behav Sci 5:100–107.

Khurana A, Romer D, Betancourt LM, Brodsky NL, Giannetta JM, Hurt H (2012) Early adolescent sexual debut: The mediating role of working memory ability, sensation seeking, and impulsivity. Dev Psychol 48:1416–1428.

Kirby KN, Maraković NN (1995) Modeling myopic decisions: Evidence for hyperbolic delay-discounting within subjects and amounts. Organ Behav Hum Dec 64:22–30.

Klein A, Ghosh SS, Avants B, Yeo BT, Fischl B, Ardekani B, Gee JC, Mann JJ, Parsey RV (2010) Evaluation of volume-based and surface-based brain image registration methods. Neuroimage 51:214–220.

Lenroot RK, Gogtay N, Greenstein DK, Wells EM, Wallace GL, Clasen LS, Blumenthal JD, Lerch J, Zijdenbos AP, Evans AC, Thompson PM, Giedd JN (2007) Sexual dimorphism of brain developmental trajectories during childhood and adolescence. Neuroimage 36:1065–1073.

Mackey, AP, Finn AS, Leonard JA, Jacoby-Senghor DS, West MR, Gabrieli CF, Gabrieli JD (2015) Neuroanatomical correlates of the income-achievement gap. Psychol Sci 26:925–933.

Marcus D, Wang T, Parker J, Csernansky JG, Morris JC, Buckner R (2007) Open Access Series of Imaging Studies (OASIS): cross-sectional MRI data in young, middle aged, nondemented, and demented older adults. J Cogn Neurosci 19:1498–1507.

Mazur JE (1987) An adjusting procedure for studying delayed reinforcement. Commons, ML.; Mazur, JE.; Nevin, JA, 55–73.

Moore TM, Reise SP, Gur RE, Hakonarson H, Gur RC (2015) Psychometric properties of the Penn Computerized Neurocognitive Battery. Neuropsychology 29:235–246.

Olson EA, Collins PF, Hooper CJ, Muetzel R, Lim KO, Luciana M (2009) White matter integrity predicts delay discounting behavior in 9-to 23-year-olds: a diffusion tensor imaging study. J Cogn Neurosci 21:1406–1421.

Peters J, Büchel C (2011) The neural mechanisms of inter-temporal decision-making: understanding variability. Trends Cogn Sci 15:227–239.

Reynolds B (2004) Do high rates of cigarette consumption increase delay discounting?: A cross-sectional comparison of adolescent smokers and young-adult smokers and nonsmokers. Behav Process 67:545–549.

Romer D, Reyna VF, Satterthwaite TD (2017) Beyond stereotypes of adolescent risk taking: placing the adolescent brain in developmental context. Dev Cogn Neurosci 27:19–34.

Rosen A, Roalf DR, Ruparel K, Blake J, Seelaus K, Villa LP, Ciric R, Cook PA, Davatzikos C, Elliott MA, Garcia De La Garza A, Gennatas ED, Quarmley M, Schmitt JE, Shionhara RT, Tisdall MD, Craddock RC, Gur RE, Gur RC, Satterthwaite TD (2017) Data-driven assessment of structural image quality. bioRxiv doi: 10.1101/123456.

Satterthwaite TD, Elliott MA, Ruparel K, Loughead J, Prabhakaran K, Calkins ME, Hopson R, Jackson C, Keefe J, Riley M, Mentch FD, Sleiman P, Verma R, Davatzikos C, Hakonarson H, Gur RC, Gur RE (2014a) Neuroimaging of the Philadelphia neurodevelopmental cohort. Neuroimage 86:544–553.

Satterthwaite TD, Shinohara RT, Wolf DH, Hopson RD, Elliott MA, Vandekar SN, Ruparel K, Calkins ME, Roalf DR, Gennatas ED, Jackson C, Erus G, Prabhakaran K, Davatzikos C, Detre JA, Hakonarson H, Gur RC, Gur RE (2014b) Impact of puberty on the evolution of cerebral perfusion during adolescence. Proc Natl Acad Sci U S A 111:8643–8648.

Satterthwaite TD, Connolly JJ, Ruparel K, Calkins ME, Jackson C, Elliott MA, Roalf DR, Hopson R, Prabhakaran K, Behr M, Qiu H, Mentch FD, Chiavacci R, Sleiman PMA, Gur RC, Hakonarson H, Gur RE (2016) The Philadelphia Neurodevelopmental Cohort: a publicly available resource for the study of normal and abnormal brain development in youth. Neuroimage 124:1115–1119.

Scheres A, Dijkstra M, Ainslie E, Balkan J, Reynolds B, Sonuga-Barke E, Castellanos FX (2006) Temporal and probabilistic discounting of rewards in children and adolescents: effects of age and ADHD symptoms. Neuropsychologia 44:2092–2103.

Schwartz DL, Mitchell AD, Lahna DL, Luber HS, Huckans MS, Mitchell SH, Hoffman WF (2010) Global and local morphometric differences in recently abstinent methamphetamine-dependent individuals. Neuroimage 50:1392–1401.

Seeley WW, Menon V, Schatzberg AF, Keller J, Glover GH, Kenna H, Reiss AL, Greicius MD (2007) Dissociable intrinsic connectivity networks for salience processing and executive control. J Neurosci 27:2349–2356.

Sellitto M, Ciaramelli E, di Pellegrino G (2010) Myopic discounting of future rewards after medial orbitofrontal damage in humans. J Neurosci 30:16429–16436.

Senecal N, Wang T, Thompson E, Kable JW (2012) Normative arguments from experts and peers reduce delay discounting. Judgm Decis Mak 7:568–589.

Shamosh NA, Gray JR (2008) Delay discounting and intelligence: A meta-analysis. Intelligence 36:289–305.

Shamosh NA, DeYoung CG, Green AE, Reis DL, Johnson MR, Conway AR, Engle RW, Braver TS, Gray JR (2008) Individual differences in delay discounting: relation to intelligence, working memory, and anterior prefrontal cortex. Psychol Sci 19:904–911.

Sotiras A, Resnick SM, Davatzikos C (2015) Finding imaging patterns of structural covariance via non-negative matrix factorization. Neuroimage 108:1–16.

Sotiras A, Toledo JB, Gur RE, Gur RC, Satterthwaite TD, Davatzikos C (2017) Patterns of coordinated cortical remodeling during adolescence and their associations with functional specialization and evolutionary expansion. Proc Natl Acad Sci U S A 114:3527–3532.

Soutschek A, Ruff CC, Strombach T, Kalenscher T, Tobler PN (2016) Brain stimulation reveals crucial role of overcoming self-centeredness in self-control. Sci Adv 2:e1600992.

Sowell ER, Thompson PM, Leonard CM, Welcome SE, Kan E, Toga AW (2004) Longitudinal mapping of cortical thickness and brain growth in normal children. J Neurosci 24:8223–8231.

Stanger C, Elton A, Ryan SR, James GA, Budney AJ, Kilts CD (2013) Neuroeconomics and adolescent substance abuse: individual differences in neural networks and delay discounting. J Am Acad Child Adolesc Psy 52:747–755.

Steinberg L, Graham S, O’Brien L, Woolard J, Cauffman E, Banich M (2009) Age differences in future orientation and delay discounting. Child Dev 80:28–44.

Strombach T, Weber B, Hangebrauk Z, Kenning P, Karipidis II, Tobler PN, Kalenscher, T (2015) Social discounting involves modulation of neural value signals by temporoparietal junction. Proc Natl Acad Sci U S A 112:1619–1624.

Thomason ME, Dassanayake MT, Shen S, Katkuri Y, Alexis M, Anderson AL, Yeo L, Mody S, Hernandez-Andrade E, Hassan SS, Studholme C, Jeong J, Romero R (2013) Cross-hemispheric functional connectivity in the human fetal brain. Sci Transl Med 5:173ra24.

Tjur T (2009) Coefficients of determination in logistic regression models—A new proposal: The coefficient of discrimination. Am Stat 63:366–372.

Tustison NJ, Avants BB, Cook PA, Zheng Y, Egan A, Yushkevich PA, Gee JC (2010) N4ITK: improved N3 bias correction. IEEE Trans Med Imaging 29:1310–1320.

Tustison NJ, Cook PA, Klein A, Song G, Das, SR, Duda JT, Kandel BM, van Strien N, Stone JR, Gee JC, Avants BB (2014) Large-scale evaluation of ANTs and FreeSurfer cortical thickness measurements. Neuroimage 99:166–179.

Vandekar SN, Shinohara RT, Raznahan A, Roalf DR, Ross M, DeLeo N, Ruparel K, Verma R, Wolf DH, Gur RC, Gur RE (2015) Topologically dissociable patterns of development of the human cerebral cortex. J Neurosci 35:599–609

Van Essen DC (2005) A population-average, landmark-and surface-based (PALS) atlas of human cerebral cortex. Neuroimage 28:635–662.

Van Essen DC, Drury HA, Dickson J, Harwell J, Hanlon D, Anderson CH (2001) An integrated software suite for surface-based analyses of cerebral cortex. J Am Med Inform Assoc 8:443–459.

Van Leijenhorst L, Moor BG, de Macks, ZAO, Rombouts SA, Westenberg PM, Crone EA (2010) Adolescent risky decision-making: neurocognitive development of reward and control regions. Neuroimage 51:345–355.

Wang H, Suh JW, Das SR, Pluta J, Craige C, Yushkevich PA (2012) Multi-atlas segmentation with joint label fusion. IEEE Trans Pattern Anal Mach Intell 35:611–623.

Wang Q, Chen C, Cai Y, Li S, Zhao X, Zheng L, Zhang H, Liu J, Chen C, Xue G (2016) Dissociated neural substrates underlying impulsive choice and impulsive action. Neuroimage 134:540–549.

Wood SN (2004) Stable and efficient multiple smoothing parameter estimation for generalized additive models. J Am Stat Assoc 99:673–686.

Wood SN (2011) Fast stable restricted maximum likelihood and marginal likelihood estimation of semiparametric generalized linear models. J Roy Stat Soc Series B Stat Meth 73:3–36.

Zielinski BA, Gennatas ED, Zhou J, Seeley WW (2010) Network-level structural covariance in the developing brain. Proc Natl Acad Sci U S A 107:18191–18196.

